# The legacy of past heatwaves on ‘off-host’ parasite stages: Reduced infection risk and costs in the *Daphnia-Pasteuria* system

**DOI:** 10.1101/2025.09.10.675284

**Authors:** Justine Boutry, Noa Lavi Shasha, Frida Ben-Ami

## Abstract

Heatwaves challenge our understanding of how environmental stressors reshape host-parasite interactions, particularly during the critical “off-host” stages of endoparasites. These stages, often dismissed as inert and unaffected by environmental changes, remain a black box in ecological research despite their key role in parasites’ life cycle. Here, we examined how heatwaves affect the infectivity and progression of infection of Pasteuria ramosa, a bacterial parasite of the planktonic crustacean Daphnia magna. In order to measure the respective influences of the bacterial genotype and the heatwaves, we exposed the off-host stages (=spores) from four genetically distinct parasite clones to three temperature regimes (20°C, 30°C, 40°C), and tested their subsequent effects on host and parasite traits. Heatwaves reduced parasite infectivity, regardless of parasite genotype. Intriguingly, heat-stressed spores induced genotype-specific shifts in their host: one host clone increased castration rates at higher temperatures, while others experienced moderate to dramatic declines. Furthermore, individuals exposed to heat-stressed spores that remained uninfected, exhibited enhanced survival, suggesting reduced costs of resistance against damaged parasite spores. This study provides insights into how heatwaves, endured only by the parasite, can modulate various aspects of host-parasite interactions, underlying the importance of the parasite’s environmental history in evolutionary ecology and epidemiology.

**Open Research Statement:** All data and code supporting the results of this study are provided here: https://github.com/JustineBoutry/HeatwaveSpores

## Introduction

With the acceleration of global climate change, understanding the evolutionary responses of parasites and their impact on epidemics and ecosystems is becoming increasingly critical (King, Hall, and Wolinska 2023). Predicting parasite dynamics under environmental fluctuations remains challenging due to our poor understanding of how external stressors reshape inherently complex host-parasite interactions (Lõhmus and Björklund 2015). Unlike ectoparasites, which are directly exposed to environmental changes and thus more easily studied, endoparasites reside within their hosts, which may offer some degree of protection from external climate fluctuations. Consequently, some authors have argued that endoparasites are mainly subjected to indirect effects of environmental fluctuations via their host (Cizauskas et al. 2017). Little is known about the direct impact of environmental fluctuations on the parasite’s life cycle, particularly during the stages spent outside the host (Marcus et al. 2023; Shodipo et al. 2020).

The impact of environmental fluctuations on endoparasites is often studied via intra-host dynamics, but this approach often overlooks the parasite’s life cycle. Indeed, many parasites spend significant portions of their life outside their host, and are directly affected by the environment. These environmentally-transmissible parasite stages can persist for extended periods—from days to centuries as evidenced, for example, by viruses retained in permafrost while maintaining infectivity (Rigou et al. 2022) and microsporidia and bacterial pathogens surviving decades in lake sediments (Decaestecker et al. 2004)—yet the biological and ecological implications of climate change on these “off-host” stages remain understudied. As the likelihood of extreme events, such as droughts and heatwaves, is expected to increase in the near future, understanding how parasites respond to such events and how these responses affect ecosystems has become urgent (Pörtner et al. 2022).

Parasites can induce profound ecological shifts when affecting keystone species, particularly in aquatic ecosystems (Lafferty et al. 2008; Thomas et al. 1998). In this perspective, the planktonic crustacean *Daphnia magna* serves as an exemplary model system for investigating host-parasite dynamics in ecological and evolutionary contexts (Ebert 2022). When infected by the endoparasitic bacterium *Pasteuria ramosa* that induces host castration (Ebert et al. 2004), dramatic alterations of *Daphnia* population demographics and communities occur (Goren and Ben-Ami 2013; Halle et al. 2024). Moreover, its prolonged environmental persistence through spore formation enables extended transmission capabilities (Decaestecker et al. 2004). Spore-forming bacteria have gained attention due to their exceptional environmental resilience and capacity for long-term transmission, with potential for substantial ecological perturbations (Setlow 2014). The ability of *P. ramosa* to persist in sediments parallels other spore-forming human pathogens, such as *Bacillus anthracis* and *Clostridium tetani* (Carlson et al. 2018; Anellis, Grecz, and Berkowitz 1965), thus underscoring the epidemiological significance of this model system.

Despite evidence that heatwaves can reduce the infectivity of *P. ramosa* (Marcus et al. 2023), the underlying mechanisms remain unclear. This study investigated how genotype-specific responses to heatwaves during the off-host stage shape parasite fitness. By implementing a comprehensive genotype-by-environment (G×E) experimental design, we quantitatively assessed how heatwaves endured only by the parasite modulate host and parasite traits. Our experimental framework evaluated whether heatwaves function as a selective filter, potentially selecting for heat-tolerant genetic variants, or uniformly compromise spore viability across genetic backgrounds. Specifically, we tested the hypothesis whether heatwaves selectively eliminated heat-sensitive parasite genotypes or alternatively, if heatwaves uniformly reduced the infectivity of spores independently of their genotypes. We also explored how these heat-stressed spores impacted host survival and reproduction, in both infected and exposed but uninfected hosts. By linking parasite environmental history to infection phenotypes and host outcomes, our work highlights how extreme climatic events influence parasite resilience and host-parasite dynamics through dormant stages environmental legacy.

## Material and methods

### 1. Biological model

All *Daphnia magna* individuals used in this experiment belong to clone HU-HO-2 that originated in Bogarzoto, Hungary. This allowed us to measure only the effects of heatwaves on *P. ramosa* infections, avoiding genetic interactions between the *Daphnia* host and the *Pasteuria* parasite clones (Luijckx et al. 2011). Throughout our experiment, *Daphnia* were kept in ADaM (Artificial *Daphnia* Medium), prepared according to methods described in (Klüttgen et al. 1994). *Daphnia* cultures (F0) from which we sampled our individuals were previously fed 5 times per week under a 12:12 light regime, at 20°C, for 50 days. All *Daphnia* were fed 5 to 7 times per week with increasing amounts of the unicellular algae *Scenedesmus gracilis* cells as follows: 1–3 days old: 1×10^6^, 4–8 days old: 2×10^6^, 9–14 days old: 3×10^6^, 15– 85 17 days old: 5×10^6^, 18–21 days old: 6×10^6^, 22–26 days old: 7×10^6^, 27–29 days old: 8×10^6^, 30–36 days old: 9×10^6^ and greater than 36 days old: 10×10^6^.

### 2. Parasite preparation

In this study we used lab stocks (frozen dead infected individuals) of four isogenic genotypes (clones) of the semelparous bacterium *Pasteuria ramosa* from different origins (C1 from Moscow, Russia, C14 and C18 from Tvärminne, Finland and C24 from Heverlee, Belgium; see Luijckx et al. 2011). The *Daphnia* carcasses were merged according to their strain in three Eppendorf tubes. After removing as much medium (ADaM) as possible using a Pasteur pipette, the Eppendorf tubes were left open for 24 hours to dry completely. Each of these tubes was subjected to a heatwave for 72 hours using a heating block at 20°C (control without heatwave), 30°C or 40 °C. Afterwards, the *Daphnia* carcasses were crushed in the Eppendorf tubes using pestles in 100 µl of deionized water. The presence or absence of other visible stages of the parasite (e.g., Cauliflower, Grape seed, Immature, see supplementary figure S2A) were also recorded for each sample (see experimental details in supplementary materials). Mature spore concentration of each parasite clone was determined using a Thoma cell counting chamber according to the formula: Concentration (cells.mL^-1^) = 400 × Number of cells × 20 × 10^6^. All individuals were infected on the same day in April 2024 and the overall experiment ended in November 2024.

### 3. Infection

In each treatment, between 29 to 49 (Table S1) five days old *Daphnia* were each exposed to 100,000 spores. In addition, 31 *Daphnia* were exposed to a solution of crushed uninfected *Daphnia* (Table S1), resulting in a total number of 490 *Daphnia* across 13 treatments (four parasite clones × three temperatures + uninfected controls). During the experimental infection process, *Daphnia* were kept in individual jars with 20 ml of ADaM and split in 50 trays with one individual of each treatment in each tray (or batch). After 5 days we transferred each individual to a new jar with 80 ml of ADaM. For the entire duration of the experiment, *Daphnia* were kept in a climate-controlled room at 20°C with a 12:12 light cycle.

### 4. Experimental measurements

We monitored host survival and offspring production every three to four days. Dead *Daphnia* were collected and stored without medium at −20°C for later spore counting. Twice a week, we removed offspring and measured the timing of reproduction, and the number of clutches and offspring released. By day 67, most infected *Daphnia* had already died, so we ended the experiment by collecting the remaining *Daphnia* and storing them at −20°C as well. Each *Daphnia* was crushed in order to observe the infectious status according to the presence or absence of at least one parasite stage.

### 5. Statistical methods

*Daphnia* that had died before day 15 were excluded from the analysis, as their death was most likely not related to infection at this stage of the experiment. Nine individuals were also excluded because of accidental loss. Thus, the final dataset included 381 individuals in total (Table S1). Generalized linear models (GLMs) were fitted using the glmmTMB package in R (R version 4.3.1 and RStudio version 2023.12.1+402). We selected the best fitted models using the multistep approach described in (Zuur et al. 2009). More precisely, we first evaluated the potential impact of the batch as a random effect followed by the inclusion of various combinations of fixed effects (presented in the corresponding supplementary tables). Models were compared using AICc (conditional Akaike criterion) and only the most relevant effects were kept in the final model. Finally, we checked the model residuals with the package DHARMa to ensure assumptions were met (including dispersion), thereby confirming the adequacy of the final model. For survival models, we tested the proportional hazards assumption for fixed effects using Schoenfeld residuals, ignoring the random effect structure (after checking that both types of models reached the same conclusions).

Infection probability was analyzed using binary logistic regression (infected vs. uninfected). For survival analysis, we used the packages coxme and survival to construct different Cox regression models for infected and uninfected *Daphnia*. For the uninfected individuals, we initially fitted a Cox proportional hazards model including temperature as a categorical predictor. As the proportional hazards (PH) assumption was moderately violated (global test: p = 0.035), we also fitted an extended model including time-dependent covariates via interactions between temperature and log-transformed, centered survival time. This model successfully corrected the PH violation (global test: p = 0.30) and produced effect estimates consistent with the simpler model. As both models led to similar biological interpretations, we retained the standard Cox model for clarity. Full code and results of the time-dependent model are provided in the online supplementary materials.

Host reproduction was analyzed using two approaches, by either focusing on the risk of being fully castrated by the parasite (i.e., individuals that never reproduced) using a binomial distribution, or considering the number of offspring produced for individuals that produced at least one clutch, using a Gaussian distribution (for uninfected individuals), as it provided a better fit to the residual structure and allowed full data inclusion, and a negative binomial distribution (for infected individuals), to account for overdispersion of count data in offspring number. All figure designs and data visualizations were created using ggplot2.

## Results

### 1. Parasite infectivity

Heatwaves reduced the infectivity of the parasite (Figure 1A; Odds Ratio [OR]: 0.45, 95% CI: 0.31–0.65). Spores treated with 40°C heatwaves had only about half of the infectivity of control spores (Figure 1A). Relevant models accounted for the random effect of the experimental batch, but no significant effect of bacterial genotype was observed (Table S2).

**Figure 1.**
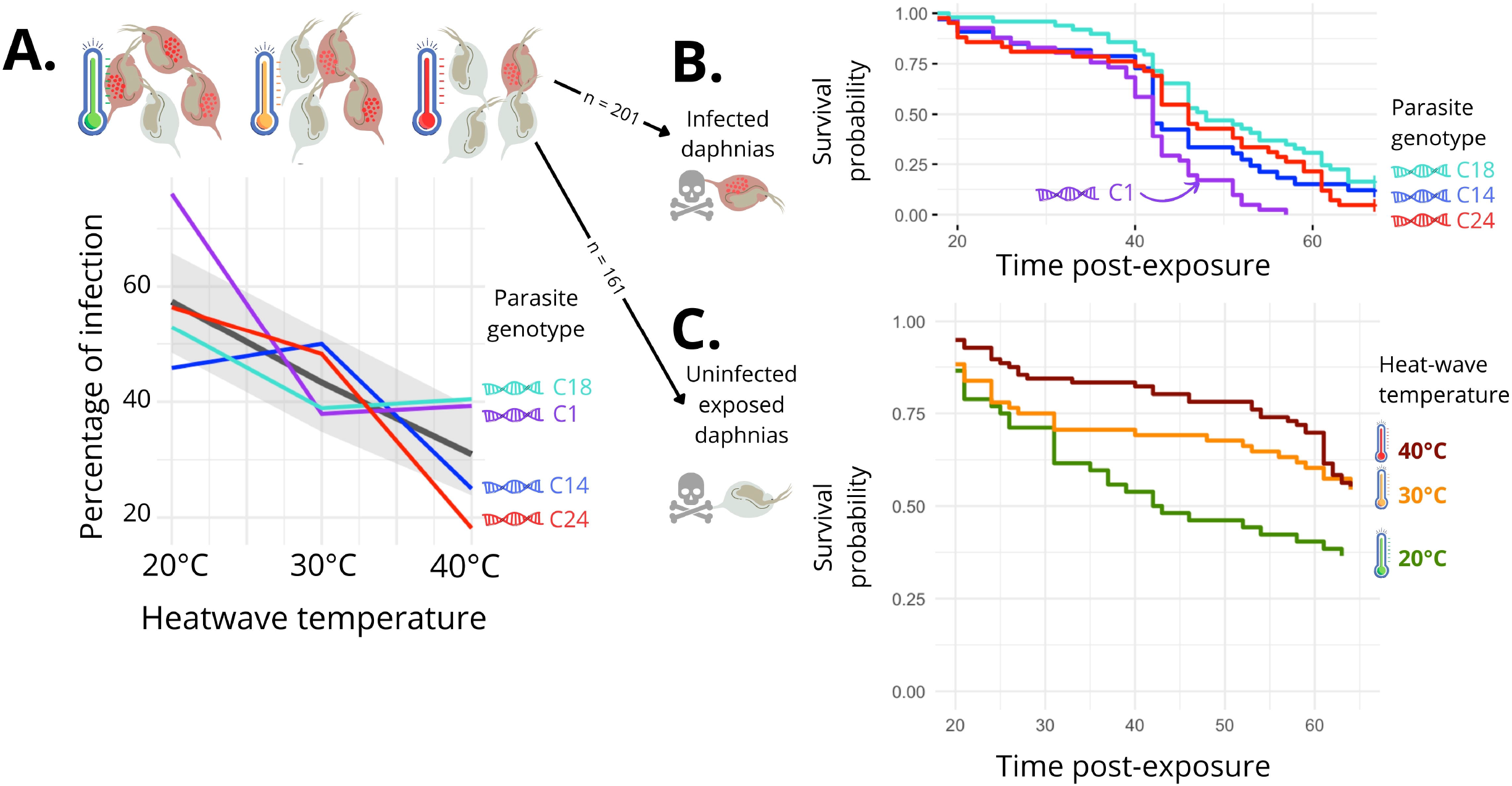
Effects of heatwave temperature and parasite genotype on infection rates and survival of *Daphnia* hosts. (A) Percentage of infected *Daphnia* hosts exposed to four parasite genotypes (C1 in purple, C14 in blue, C18 in green, and C24 in red) across three heatwave temperatures (20°C, 30°C, and 40°C). The black line represents the decrease in the average infection rate with increasing temperature. (B) Survival probability over time of *Daphnia* hosts infected with four parasite genotypes (same colours as in A). (C) Survival probability over time of exposed but uninfected *Daphnia* hosts increased with heatwave temperatures (20°C in green, 30°C in orange, and 40°C in red). Figure created by Justine Boutry using icons (Daphnia, DNA, thermometer, and skull and crossbones) from Canva Pro under a license that permits commercial reuse. Icons sourced from Canva Pro: https://www.canva.com/policies/content-license-agreement/

### 2. Host mortality

Individuals infected with spores of the C1 genotype suffered higher mortality rates than other genotypes (Figure 1B; Hazard Ratio [HR]: 1.90, 95% CI: 1.12–3.23), insofar that their virulence was nearly doubled compared to the other genotypes. The random effect for batch was significant, but temperature effects were not included in the most relevant model (Table S3).

In the case of exposed individuals that did not become infected, spores treated at a higher temperature increased the host’s survival probability by 37% (Figure 1C; HR: 0.64, 95% CI: 0.47–0.88). Neither spore genotype nor random batch effects were significant in these models (Table S4).

### 3. Host reproduction

One of the hallmark traits of *P. ramosa* infections is the parasite’s ability to castrate its host. This ability of the parasite was reduced with increasing temperatures (Figure 2A; OR: 0.11, 95% CI: 0.02–0.75), and strongly depended on bacterial genotype. While genotypes C18 and C24 experienced an overall increase in the proportion of infected individuals producing at least one clutch at higher temperatures, genotypes C14 and C1 showed an opposite trend, with the proportion of individuals producing at least one clutch nearly doubling from 20°C to 40°C for C14, and decreasing six-fold for C1 (Figure 2A; OR: 67.39, 95% CI: 4.75–955.45). No random batch effects were relevant (Table S5).

**Figure 2.**
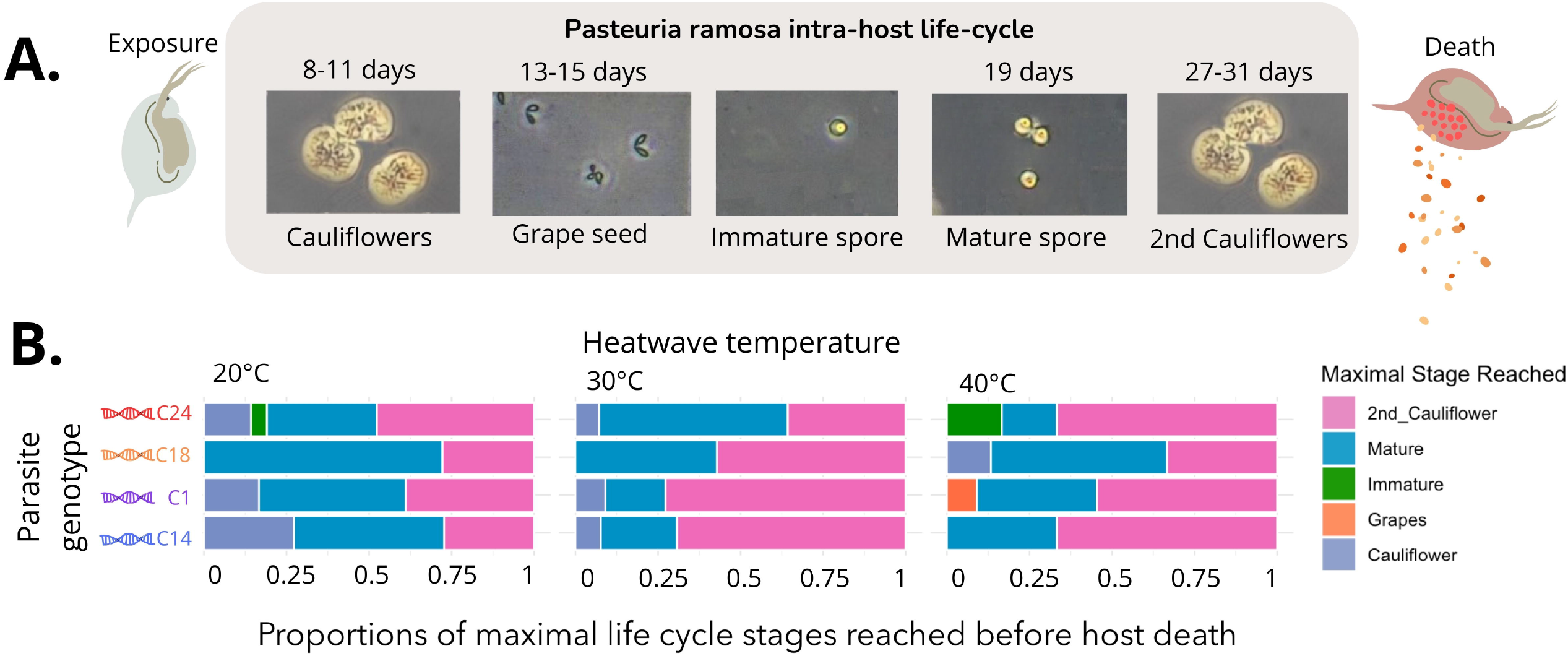
Impact of heatwaves on reproduction in infected and exposed but uninfected hosts exposed to *Pasteuria ramosa* spores. (A) Ratio of fully castrated infected *Daphnia* across different heatwave temperatures (20°C, 30°C, 40°C) and parasite genotypes (C1: purple, C14: blue, C18: green, C24: red). Bars represent mean ratios with error bars indicating standard deviations. (B) Number of offspring produced by infected individuals over time post-exposure at different temperatures. Data points represent observed values for the different genotypes (same colours as in A) and the horizontal green dashed bar represents the average cumulative number of offspring produced by exposed but uninfected *Daphnia*. Figure created by Justine Boutry using icons (Daphnia, DNA, thermometer) from Canva Pro under a license that permits commercial reuse. Icons sourced from Canva Pro: https://www.canva.com/policies/content-license-agreement/

For exposed hosts that did not become infected, only survival time influenced their reproduction, thereby increasing their chance to produce at least one clutch (OR: 0.88, 95% CI: 0.82–0.94, see supplementary figure S1A). A significant random batch effect was observed for survival time (Table S6).

For infected hosts that still produced at least one clutch, their reproduction was reduced when infected by genotypes C1 (Figure 2B; OR: 0.44, 95% CI: 0.22–0.89) and C18 (Figure 2B; OR: 0.48, 95% CI: 0.28–0.82) in comparison to genotypes C14 and C24, regardless of temperature (Table S7). Over time, offspring production of all infected individuals increased by 3% every day, while offspring production of exposed hosts that did not become infected increased by 30% every day (estimate = 1.30, 95% CI: 1.16–1.45, see supplementary figure S1B). No random batch effects were significant (Table S8).

## Discussion

Understanding how climate extremes affect parasite transmission remains a key challenge in disease ecology. Here we show that heatwaves can reduce parasite infectivity by acting on parasites during their environmental stages, with consequences for host-parasite dynamics. Contrary to the hypothesis that heatwaves should select for heat-tolerant genotypes, our findings suggest that heatwaves act as a uniform stressor, reducing spore infectivity regardless of their genotype. This challenges the prediction that heat stress directly drives the selection and evolution of heat-resistant variants, emphasizing the importance of considering environmentally-induced plasticity in understanding parasite resilience. These thermal effects extended beyond infected individuals, altering outcomes even for exposed but uninfected hosts, resulting in improved survival compared to those exposed to non-stressed spores. An additional layer of complexity emerged in how heatwaves altered parasite-induced castration, depending on the heatwave temperature, but differently for each parasite genotype. These findings suggest that heatwaves not only affect parasite infectivity, but also modulate host-parasite interactions and their broader ecological implications.

### Insights into the effects of heatwaves on parasite infectivity

Our findings further challenge the prediction that spore survival through heatwaves would be mediated by genetically encoded heat-resistance, similar to global resistance mechanisms to infection (Bento et al. 2017; Luijckx 2012). While supporting previous findings (Marcus et al. 2023), our results show that high temperatures reduced spore infectivity across all genotypes, highlighting plastic responses over genetic resistance. This advocates for the importance of environmental effects and plastic responses of the parasite in shaping infection dynamics (Vale et al. 2011). Of course, this does not rule out other evolutionary mechanisms, as untested genotypes may exhibit distinct responses.

### Reduced cost of parasite defenses and immunological implications

Heat-stressed spores exhibited reduced infectivity and thus may have imposed a lower cost of defense on exposed but uninfected hosts, as indicated by their improved survival compared to animals exposed to non-stressed spores. This surprising result suggests that damaged spores are less effective at triggering host defenses, resulting in lower costs to the host and increased host survival post-exposure (see supplementary materials for detailed analysis of effects across parasite life-cycle stages). Furthermore, this specific host defense appears to be independent of the spores genotype, in contrast to other resistance mechanisms that have a genetic basis (Luijckx et al. 2011; Bento et al. 2017; Ebert et al. 2016). Instead, host defense is principally influenced by previous heatwaves experienced by the parasite. If such effects prove generalizable to other parasite systems and operate at the ecosystem scale, reduced immune costs following a heatwave and exposure to environmentally compromised parasites could profoundly alter the outcome host–parasite interactions and the structure of ecological communities (Hall et al. 2024; Brooks and Hoberg 2007).

### Environmental modulation of virulence and host manipulation

A particularly compelling aspects of our study is the impact heatwaves had on parasite-induced host castration, a well-documented strategy of parasites to divert host resources towards their own growth and reproduction (Ebert et al. 2004; Poulin 2010). In our study, parasite virulence, measured as parasite-induced host mortality, did not appear to be affected by heatwaves and remained primarily genotype-dependent (see also Clerc, Ebert, and Hall 2015). However, the proportion of individuals that never reproduced (total castration) varied across parasite genotypes, after parasite spores had been exposed to heatwaves. For example, C14, the genotype that caused the highest proportion of total castration under moderate heatwaves, exhibited a sharp decline in total castration ability at 40°C. Conversely, C1, which showed average total castration abilities under moderate heatwaves, castrated nearly all hosts at 40°C. These responses suggest that different parasite genotypes employ distinct mechanisms to achieve host castration, possibly relying on plastic responses. The adaptive value of such modulations remains to be determined, as it does not seem to be associated with increased or decreased parasite fitness (see supplementary analysis on parasite life-cycle, figure S3 and S4).

### Future directions and ecological relevance

Our study reveals that extreme thermal events can significantly disrupt parasite transmission by acting directly on free-living stages outside the host. This disruption reduces parasite infectivity and alters host-parasite dynamics in ways that are likely to affect other species and systems as well. Many parasites, including spore-forming bacteria and helminths, rely on environmental stages to persist or transmit (Walther and Ewald 2004; Carlson et al. 2018; Smits et al. 2016), suggesting that climate extremes can broadly interfere with these critical phases.

Beyond the immediate effects on infection, our results invite broader reflection on the ecological and evolutionary consequences of abiotic stress. In systems where transmission depends on environmental persistence, thermal fluctuations may reshape not only parasite success, but also the structure of ecological networks through changes in host susceptibility, parasite prevalence, and indirect interactions. Given that parasites often play key roles in regulating food webs, influencing species coexistence, and modulating energy flow (Brooks and Hoberg 2007; Thomas et al. 1999), disruptions occurring at the transmission stage could have far-reaching implications. Recognizing parasites as integral ecological actors underscores the need to incorporate their full range of responses to environmental variability into ecosystem models (Brooks and Hoberg 2007).

These dynamics also resonate with One Health perspectives, which emphasize the interconnectedness of environmental, animal, and human health (Panel (OHHLEP) et al. 2022). In our system, parasites exposed to environmental stress influenced infection outcomes in hosts that had not themselves experienced the disturbance. Notably, even the survival of uninfected hosts was modulated by the heatwaves endured by spores, suggesting that parasite exposure to abiotic stress can alter host outcomes independently of infection. This non-intuitive effect illustrates how environmental stressors acting on parasites can cascade through host populations, including those not directly exposed, thereby influencing broader ecological and epidemiological dynamics. Such cross-scale effects reinforce the need to integrate ecological, epidemiological, and environmental frameworks when predicting the consequences of climate variability on disease systems (Ghai et al. 2022).

Our findings highlight the need to broaden current perspectives of parasite ecology under climate change. While rising temperatures can be conceptualized as a long-term selective filter favoring more climate-resilient strains, our results show that short-term thermal extremes can have immediate, genotype-independent effects on parasite performance. These effects are particularly critical when parasites are directly exposed to environmental stress during free-living or dormant stages, outside the host. Such disruptions can modulate infection outcomes and reshape host-parasite interactions before infection even occurs. Accounting for these overlooked stages is essential for anticipating disease emergence and forecasting how global change will reshape ecological interactions.

## Acknowledgments

J.B. was supported by a fellowship from the Azrieli Foundation and by funding from the George S. Wise Faculty of Life Sciences at Tel Aviv University. We thank Enav Bereza for maintaining the laboratory algae, and all lab members who contributed to the early stages of the experiment. We are grateful to both funding institutions for their support.

## Author Contributions

J.B. and F.B.A. conceived and designed the study, J.B. conducted all experiments and analyses, and drafted the manuscript. N.L.S. assisted with experiment maintenance and data collection. F.B.A. secured funding and provided feedback on the manuscript. All authors approved the final version of the manuscript.

---

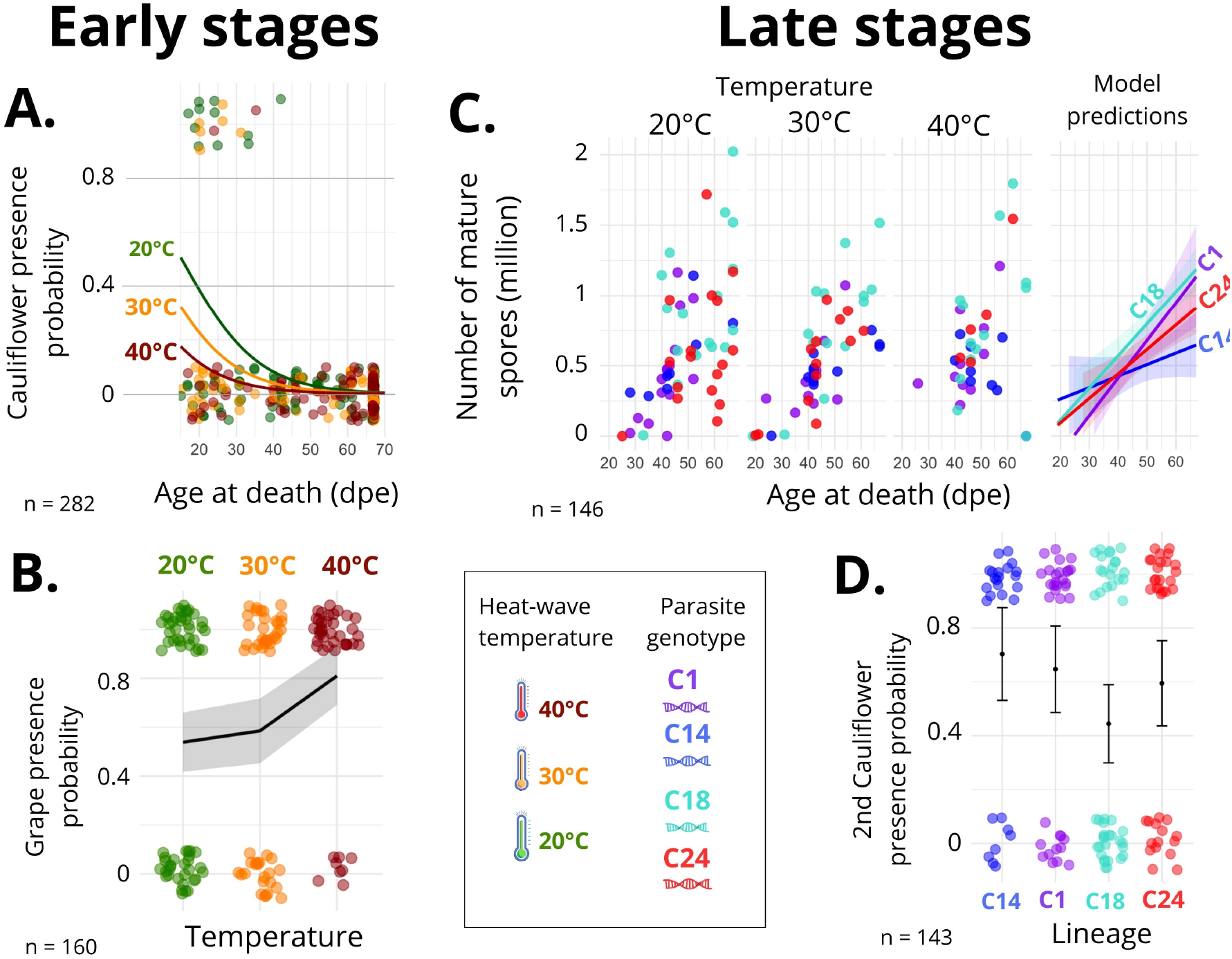

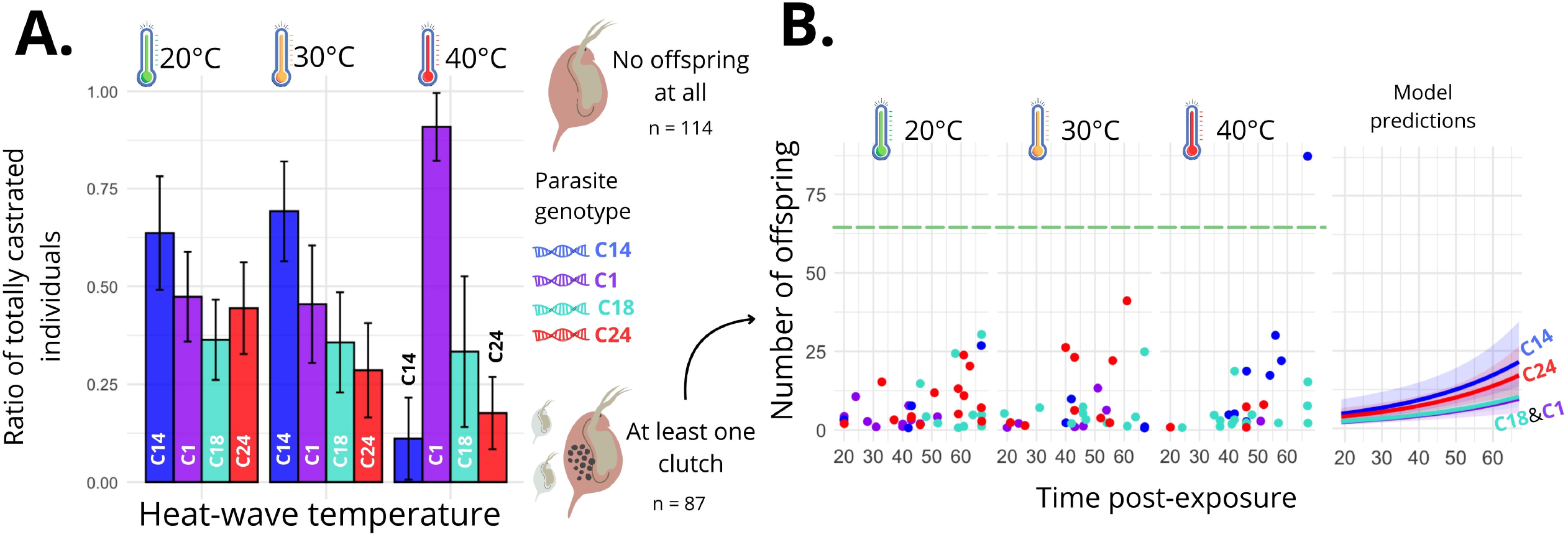

