## Supplementary material for "The legacy of past heatwaves on ‘off-host’ parasite stages: Reduced infection risk and costs in the *Daphnia-Pasteuria* system"

**Syntax Guide for Statistical Modeling**

### 1. Model Selection Process

#### Step 1: Random effect selection

- Compare models with and without (1|Batch)
- Choose based on AICc (lower = better)

#### Step 2: Fixed effect selection

- Compare different combinations of Isolate, Temperature, DeathAge
- Test additive (+) vs. interactive (*) relationships
- Choose best model based on AICc

#### Quick Reference for Our Study

| Model Formula | Biological Interpretation |
|---|---|
| Y ~ 1 + (1\|Batch) | Only random batch variation |
| Y ~ Temperature + (1\|Batch) | Temperature effect with batch variation |
| Y ~ Isolate + (1\|Batch) | Parasite genotype effect with batch variation |
| Y ~ Isolate + Temperature + (1\|Batch) | Independent effects of both factors |
| Y ~ Isolate * Temperature + (1\|Batch) | Interactive effects of both factors |
| Y ~ DeathAge + (1\|Batch) | Host longevity effect with batch variation |

### 2. General syntaxes structure

#### 2.1 Response_Variable ~ Predictors

- Left of ~: Dependent variable (what we want to predict/explain)
- Right of ~: Explanatory variables (predictors)
- The ~ symbol: Reads as "is modeled by" or "as a function of"

### 2.2 Random Effects (Mixed Models)

**Random intercept are symbolized by "(1 | Group)"**

Y ~ A + B + (1 | Batch)

*Meaning:* Intercept varies by group

*Mathematical translation:* $Y = \beta_0 + u_{0j} + \beta_1 A + \beta_2 B + \varepsilon$

- $\beta_0$: fixed intercept
- $u_{0j}$: random intercept for group j

### 2.3 Symbols Used in the formula

**Additive Effects are symbolized by "+"**

Y ~ A + B + C

*Meaning:* Y is predicted by A, B, and C independently

*Mathematical translation:* $Y = \beta_0 + \beta_1 A + \beta_2 B + \beta_3 C + \varepsilon$

**Multiplicative Effects (interactions) are symbolized by "*"**

Y ~ A * B

*Meaning:* Y is predicted by A, B, and their interaction

*Mathematical translation:* $Y = \beta_0 + \beta_1 A + \beta_2 B + \beta_3 (A \times B) + \varepsilon$

Equivalent to: Y ~ A + B + A:B

**Interaction term only are symbolized by ":"**

Y ~ A + B + A:B

*Meaning:* Main effects of A and B plus their interaction

*Mathematical translation:* $Y = \beta_0 + \beta_1 A + \beta_2 B + \beta_3 (A \times B) + \varepsilon$

**Intercept only are symbolized by "1"**

Y ~ 1

*Meaning:* Model with only an intercept (null model)

*Mathematical translation:* $Y = \beta_0 + \varepsilon$

**Terms Used in Our Tables**

| Formula Component | Interpretation | Example from Study |
|---|---|---|
| Isolate | Parasite genotype (C1, C14, C18, C24) | Fixed effect |
| Temperature | Heat treatment (20°C, 30°C, 40°C) | Fixed effect |
| DeathAge | Host age at death | Fixed effect |
| (1\|Batch) | Random intercept for experimental batch | Random effect |
| Isolate + Temperature | Independent effects of both factors | Additive model |
| Isolate * Temperature | Effects of both factors plus their interaction | Interactive model |
| 1 | Intercept-only model | Null model |

Tabe S1 : Effectives

| Heatwave treatment | C1 | | | C14 | | | C18 | | | C24 | | |
|---|---|---|---|---|---|---|---|---|---|---|---|---|
| | Included | Died<15 | Total | Included | Died<15 | Total | Included | Died<15 | Total | Included | Died<15 | Total |
| **20** | 30 | 7 | 37 | 27 | 8 | 35 | 34 | 5 | 39 | 39 | 6 | 45 |
| **30** | 29 | 5 | 34 | 27 | 7 | 34 | 37 | 7 | 44 | 28 | 1 | 29 |
| **40** | 27 | 7 | 34 | 37 | 5 | 42 | 42 | 7 | 49 | 33 | 4 | 37 |
| **Control** | 26 | 5 | 31 | | | | | | | | | |

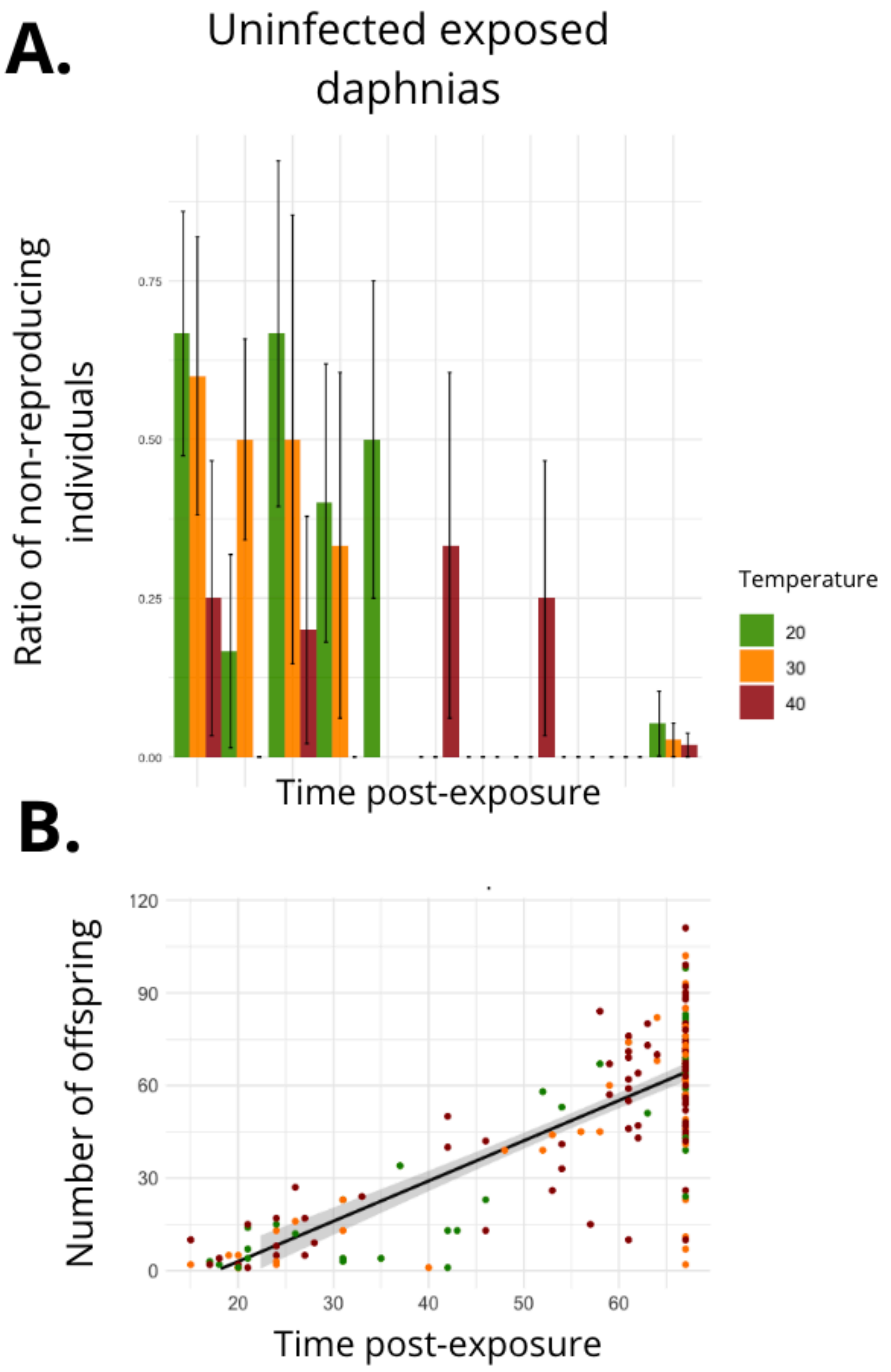


Figure S1: Effects of heatwaves endured by *Pasteuria ramosa* spores on non-infected exposed host reproduction

(A) Ratio of non-reproducing uninfected Daphnia over time post-exposure, stratified by temperature (green: 20°C, orange: 30°C, red: 40°C). Increased survival time correlated with reduced non-reproducing proportions (OR = 0.88, CI: 0.82–0.94, p = 0.0003). Bars represent means, and error bars show standard deviations.
(B) Number of offspring produced by uninfected exposed individuals increased by approximately 30% over time (OR = 1.30, CI: 1.16–1.45, $p < 2\times10^{-16}$). Data points represent observed values, with colors depending of temperatures (same as A). The solid line indicates the prediction model, with confidence intervals shaded in gray.

Table S2 : Parasite infectivity

| Step 1 : Random effect selection | |
|---|---|
| Model | AICc |
| Infected Status ~ Isolate * Temperature | 520.4 |
| Infected Status ~ Isolate * Temperature + (1 \| Batch) | 514.6 |
| Step 2 : Fixed effect selection | |
| Infected Status ~ 1 + (1 \| Batch) | 521.6 |
| Infected Status ~ Isolate + (1 \| Batch) | 525.2 |
| **Infected Status ~ Temperature + (1 \| Batch)** | **505.9** |
| Infected Status ~ Isolate + Temperature + (1 \| Batch) | 509.4 |
| InfectedStatus ~ Isolate x Temperature + (1 \| Batch) | 510.1 |

Table S3 : Mortality of infected individuals

| Step 1 : Random effect selection | |
|---|---|
| Model | AICc |
| Survival ~ Isolate * Temperature | 1306.8 |
| Survival ~ Isolate * Temperature + (1\|Batch) | 1303.8 |
| Step 2 : Fixed effect selection | |
| Survival ~ 1 + (1\|Batch) | 1306.8 |
| Survival ~ Temperature + (1\|Batch) | 1309.7 |
| **Survival ~ Isolate + (1\|Batch)** | **1293.7** |
| Survival ~ Isolate + Temperature + (1\|Batch) | 1296.7 |
| Survival ~ Isolate x Temperature + (1\|Batch) | 1303.8 |

Table S4 : Mortality of uninfected individuals

| Step 1 : Random effect selection | |
|---|---|
| Model | AICc |
| Survival ~ Isolate * Temperature | 1113.5 |
| Survival ~ Isolate * Temperature + (1\|Batch) | 1119.1 |
| Step 2 : Fixed effect selection | |
| Survival ~ 1 | 1104. |
| **Survival ~ Temperature** | **1101.7** |
| Survival ~ Isolate | 1110.6 |
| Survival ~ Isolate + Temperature | 1107.6 |
| Survival ~ Isolate x Temperature | 1113.5 |

Table S5 : Full castration of infected individuals

| Step 1 : Random effect selection | |
|---|---|
| Model | AICc |
| Never_repro~ Isolate + Temperature + DeathAge | 218.2 |
| Never_repro~ Isolate + Temperature + DeathAge + (1\|Batch) | 215.9 |
| Step 2 : Fixed effect selection | |
| Never_repro ~ DeathAge + Isolate * Temperature + (1 \| Batch) | 218.5 |
| **Never_repro ~ Isolate * Temperature + (1 \| Batch)** | **218.4** |
| Never_repro ~ DeathAge + Isolate + Temperature + (1 \| Batch) | 224.0 |
| Never_repro ~ Isolate + Temperature + (1 \| Batch) | 223.5 |
| Never_repro ~ DeathAge + Temperature + (1 \| Batch) | 222.8 |
| Never_repro ~ Temperature + (1 \| Batch) | 224.6 |
| Never_repro ~ DeathAge + Isolate + (1 \| Batch) | 220.0 |
| Never_repro ~ Isolate + (1 \| Batch) | 219.6 |
| Never_repro ~ DeathAge + (1 \| Batch) | 218.8 |
| Never_repro ~ 1 + (1 \| Batch) | 220.7 |

Table S6 : Full castration of uninfected individuals

| Step 1 : Random effect selection | |
|---|---|
| Model | AICc |
| Never_repro~ Isolate + Temperature + DeathAge | 118.5 |
| Never_repro~ Isolate + Temperature + DeathAge + (1 \| Batch) | 106.9 |
| Step 2 : Fixed effect selection | |
| Never_repro ~ DeathAge + Isolate * Temperature + (1 \| Batch) | 109.7 |
| Never_repro ~ Isolate * Temperature + (1 \| Batch) | 150.2 |
| Never_repro ~ DeathAge + Isolate + Temperature + (1 \| Batch) | 107.3 |
| Never_repro ~ Isolate + Temperature + (1 \| Batch) | 146.3 |
| Never_repro ~ DeathAge + Temperature + (1 \| Batch) | 104.0 |
| Never_repro ~ Temperature + (1 \| Batch) | 142.0 |
| Never_repro ~ DeathAge + Isolate + (1 \| Batch) | 105.0 |
| Never_repro ~ Isolate + (1 \| Batch) | 153.5 |
| ***Never_repro ~ DeathAge + (1 \| Batch)*** | ***102.8*** |
| Never_repro ~ 1 + (1 \| Batch) | 149.0 |

Table S7 : Number of offspring of reproducing infected individuals

| Step 1 : Random effect selection | |
|---|---|
| Model | AICc |
| ***Nb_offspring ~ Isolate * Temperature + DeathAge*** | ***601.9*** |
| Nb_offspring ~ Isolate * Temperature + DeathAge + (1\|Batch) | 601.8 |
| Step 2 : Fixed effect selection | |
| Nb_offspring ~ DeathAge + Isolate * Temperature | 588.0 |
| Nb_offspring ~ Isolate * Temperature | 598.8 |
| Nb_offspring ~ DeathAge + Isolate + Temperature | 585.5 |
| Nb_offspring ~ Isolate + Temperature | 598.7 |
| Nb_offspring ~ DeathAge + Temperature | 588.2 |
| Nb_offspring ~ Temperature | 607.0 |
| **Nb_offspring ~ DeathAge + Isolate** | **580.9** |
| Nb_offspring ~ Isolate | 594.1 |
| Nb_offspring ~ DeathAge | 584.3 |
| Nb_offspring ~ 1 | 604.2 |

Table S8 : Number of offspring of reproducing uninfected individuals

| Step 1 : Random effect selection | |
|---|---|
| Model | AICc |
| ***Nb_offspring ~ Isolate * Temperature + DeathAge*** | ***1649.5*** |
| Nb_offspring ~ Isolate * Temperature + DeathAge + (1\|Batch) | 1651.4 |
| Step 2 : Fixed effect selection | |
| Nb_offspring ~ DeathAge + Isolate * Temperature | 1649.5 |
| Nb_offspring ~ Isolate * Temperature | 1834.6 |
| Nb_offspring ~ DeathAge + Isolate + Temperature | 1637.8 |
| Nb_offspring ~ Isolate + Temperature | 1823.1 |
| Nb_offspring ~ DeathAge + Temperature | 1635.6 |
| Nb_offspring ~ Temperature | 1817.9 |
| Nb_offspring ~ DeathAge + Isolate | 1635.1 |
| Nb_offspring ~ Isolate | 1824.1 |
| **Nb_offspring ~ DeathAge** | **1632.8** |
| Nb_offspring ~ 1 | 1819.5 |

### Parasite life-cycle complementary analysis


#### Abstract

Hosts infected with parasite spores that endured a heatwave, harbored abnormal early-stages of the parasite (e.g., less cauliflowers and more grape formations), while late-stages of the parasite remained unaffected across all treatments, thereby ensuring stable reproductive output.


#### Material and methods

After crushing the Daphnia, for each infected sample, we recorded the presence of the different stages: cauliflower, grape seed, immature and mature spores (Dieter Ebert et al. 1996 and supplementary figure S2A).

*Statistical analyses*

Considering the different stages of the parasite's life cycle observed, in our evaluated models we also assessed the potential impact of time to timing at host death in the evaluated models, as individuals died throughout the experiment. The presence of initial cauliflowers, grape seed stages and immature spores was analyzed using a binomial distribution, while mature spore load was modelled using a Gaussian distribution. The presence of second stage cauliflowers was also evaluated using a binomial distribution. We analyzed the first and the second cycle cauliflowers separately, by splitting the dataset into two parts. We also included a global qualitative analysis of life cycle maturation, using a polynomial linear regression for modelling the maximal stage reached.

#### Results

***3. Parasite reproduction***

**3.1 Cauliflowers (first stage)**

Cauliflowers are the first visible stage of parasite maturation after esophageal penetration (Supplementary figure S2A). Heatwave temperature decreased the probability of observing cauliflowers in dead hosts (Supplementary figure S3A; OR: 0.33, 95% CI: 0.11–1.02, $p = 0.054$).

Longer host survival also reduced the likelihood of observing this stage, which is expected regarding the bacterial life-cycle. No significant effects of spore genotype or random batch effects were detected (Table S5).

**3.2 Grape seed (second stage)**

The probability of observing the grape seed stage upon host death increased with higher spore exposure temperatures (Supplementary figure S3B; OR: 2.68, 95% CI: 1.37–5.24, p = 0.004). Heat-stressed spores had a 149% higher likelihood of forming grape seed stages, regardless of host age at death. No significant effects of spore genotype or random batch effects were observed (Table S6).

**3.4 Mature spores**

Only mature spores can infect other hosts, hence, reaching maturity is vital for the parasite. Maturation speed varied among genotypes, with C1 and C18 maturing faster than C14 and C24 (Supplementary figure S3C; C1 estimate: 0.02, 95% CI: 0.00–0.04, p = 0.036; C18 estimate: 0.01, 95% CI: 0.00–0.03, p = 0.028). Temperature and random batch effects were not significant (Table S7).

**3.5 Second cycle cauliflowers**

Some mature spores, if retained long enough in their host, initiated a second reproduction cycle by producing cauliflowers. The C18 genotype was less likely to produce cauliflowers compared to other genotypes (Supplementary figure S2D; OR: 0.34, 95% CI: 0.12–0.93, p = 0.035). Neither temperature nor random batch effects were significant (Table S8).

**3.6 Maximal stage reached**

A quadratic relationship was observed between temperature and the maximal stage of maturation upon host death (Supplementary figure S3B; OR: 0.42, 95% CI: 0.23–0.76, $p$ = 0.006), indicating that maturation was more efficient at 30°C compared to 20°C and 40°C. In contrast, genotype-specific effects were marginally significant only for parasite clone C18 (OR: 0.40, 95% CI: 0.15–1.02, $p$ =

0.061), with a slightly less efficient maturation process compared to C14. No random batch effects were relevant in the models tested (Table S9).

### Discussion

#### Vulnerability of parasite life cycle after a heatwave

Our results show that heatwaves primarily affected the early stages of infection by *P. ramosa*, as indicated by the shift in the presence of its life cycle stages. Specifically, cauliflowers were less frequent, while grape seed stages were more prevalent following heatwave exposure, which suggests that heatwaves disrupt the early establishment of the parasite after host penetration. This could impair the parasite's ability to resist host defenses, which are particularly active during this phase (Ebert et al. 2016). These defenses, such as increased phenoloxidase activity (Pauwels et al. 2010), contribute to parasite clearance early in the infection process (Hall and Ebert 2012), and it has been shown that *D. magna* is capable of clearance of *P. ramosa* infections both during early parasite establishment and at progressed infection phases (Izhar et al. 2020). While early infection was sensitive to heatwaves, parasite maturation remained largely unaffected, pointing to stage-specific robustness. Indeed, neither total production of mature spores nor the presence of immature spores was affected by heatwaves. In fact, moderate heatwaves may even enhance the efficiency of the maturation process, potentially due to density-dependent growth that allows parasites to compensate for early setbacks (D. Ebert, Zschokke-Rohringer, and Carius 2000; Ben-Ami, Ebert, and Regoes 2010). This can explain why late stages of maturation are insulated from host immune responses (Little and Ebert 2000) and could provide an evolutionary advantage, by ensuring stable reproductive output in fluctuating environments.

**Figures**

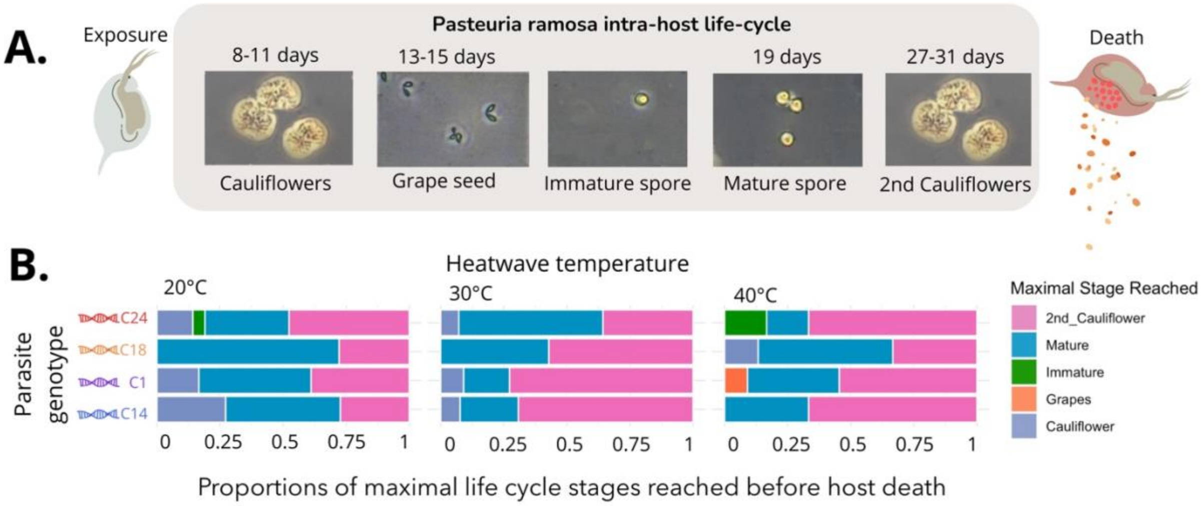


**Figure S2.**

(A) Schematic representation of the within-host life cycle of *Pasteuria ramosa* following experimental heatwaves endured by mature spores, showing progression through distinct stages: cauliflowers, grape seed stages, immature and mature spores. (B) Proportions of maximal life cycle stages reached before host death at 20°C, 30°C and 40°C. Stacked bar plots represent the proportion of hosts reaching each stage (second cauliflowers: pink, mature spores: blue, immature spores: green, grape seed stages: orange, first cauliflowers: purple). Figure created by Justine Boutry using icons (DNA) from Canva Pro under a license that permits commercial reuse. Icons sourced from Canva Pro: https://www.canva.com/policies/content-license-agreement/. Photograph by Justine Boutry, co-author and Yonathan Wiegle, used with permission (documentation uploaded separately).

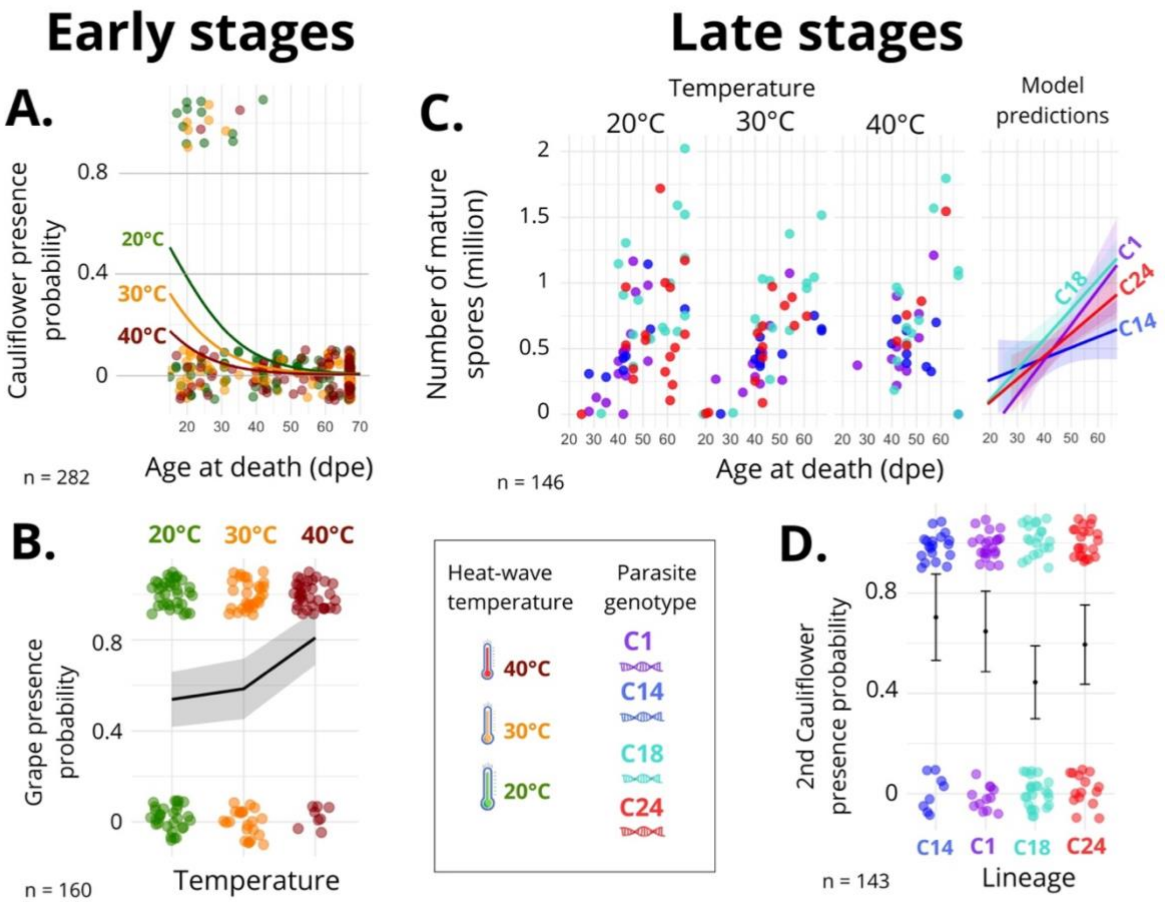


**Figure S3. Factors influencing early and late life cycle stages of *Pasteuria ramosa*.**

(A) The probability that cauliflower stages are present depending of the heatwave temperature endured by spored and the host age at death and. Dots represent data points, with model predictions represented by lines in different colours (green: 20°C, orange: 30°C, red: 40°C). (B) The probability that grape seed stages are present increased with pre-exposure heatwave temperature. Dots represent data points, with model prediction represented by a solid line with a shaded 95% CI. (C) Mature spore production by genotype. Data points represent spore counts by genotype, with model predictions shown as lines and shaded areas. (D) The probability that second-stage cauliflowers. Data points are grouped by genotype (C14: blue, C1: purple, C18: green, C24: red), with error bars representing 95% CI. Figure created by Justine Boutry using icons (DNA, Thermometers) from Canva Pro under a license that permits commercial reuse. Icons sourced from Canva Pro: https://www.canva.com/policies/content-license-agreement/.

Little, Tom J., and Dieter Ebert. 2000. "The Cause of Parasitic Infection in Natural Populations of *Daphnia* (Crustacea: Cladocera): The Role of Host Genetics." *Proceedings of the Royal*

*Society of London. Series B: Biological Sciences* 267 (1457). https://doi.org/10.1098/rspb.2000.1246.

Pauwels, Kevin, Luc De Meester, Ellen Decaestecker, and Robby Stoks. 2010. “Phenoloxidase but Not Lytic Activity Reflects Resistance against Pasteuria Ramosa in Daphnia Magna.” *Biology Letters* 7 (1): 156–59. https://doi.org/10.1098/rsbl.2010.0634.

Table S9 : Cauliflowers presence

| Step 1 : Random effect selection | |
|---|---|
| Model | AICc |
| ***cauliflowers~Temperature+Isolate+DeathAge*** | ***118.5*** |
| cauliflowers~Temperature+Isolate+DeathAge + (1\|Batch) | 120.6 |
| Step 2 : Fixed effect selection | |
| cauliflowers~Temperature*Isolate*DeathAge | NA (convergence problem) |
| cauliflowers~Temperature+Isolate*DeathAge | 119.9 |
| cauliflowers~Temperature*Isolate+DeathAge | 122.4 |
| cauliflowers~Isolate+DeathAge*Temperature | 120.0 |
| cauliflowers~Isolate*DeathAge | 120.3 |
| cauliflowers~Temperature*Isolate | 150.6 |
| cauliflowers~DeathAge*Temperature | 114.2 |
| cauliflowers~Temperature+Isolate+DeathAge | 115.9 |
| cauliflowers~Isolate+DeathAge | 116.5 |
| cauliflowers~Temperature+Isolate | 148.2 |
| ***cauliflowers~DeathAge+Temperature*** | ***110.1*** |
| cauliflowers~DeathAge | 111.2 |
| cauliflowers~Isolate | 157.0 |
| cauliflowers~Temperature | 143.7 |
| cauliflowers~1 | 152.4 |

Table S10 : Grape seed stage presence

| Step 1 : Random effect selection | |
|---|---|
| Model | AICc |
| ***Grapes~Temperature+Isolate+DeathAge*** | ***207.3*** |
| grapes~Temperature+Isolate+DeathAge +(1\|Batch) | 208.8 |
| Step 2 : Fixed effect selection | |
| grapes~Temperature*Isolate*DeathAge | 225.3 |
| grapes~Temperature+Isolate*DeathAge | 203.3 |
| grapes~Temperature*Isolate+DeathAge | 218.1 |
| grapes~Isolate+DeathAge*Temperature | 210.5 |
| grapes~Isolate*DeathAge | 206.4 |
| grapes~Temperature*Isolate | 216.0 |
| grapes~DeathAge*Temperature | 204.8 |
| grapes~Temperature+Isolate+DeathAge | 207.3 |
| grapes~Isolate+DeathAge | 212.0 |
| grapes~Temperature+Isolate | 205.6 |
| grapes~DeathAge+Temperature | 201.8 |
| grapes~DeathAge | 207.1 |
| grapes~Isolate | 210.5 |
| ***grapes~Temperature*** | ***200.1*** |
| grapes~1 | 205.7 |

Table S11 : Mature spores load

| Step 1 : Random effect selection | |
|---|---|
| Model | AICc |
| ***Mature spores load~Temperature*Isolate*DeathAge*** | ***242.1*** |
| Mature spores load~Temperature*Isolate*DeathAge + (1\|Batch) | 244.8 |
| Step 2 : Fixed effect selection | |
| Mature spores load~Temperature*Isolate*DeathAge | 242.1 |
| Mature spores load~Temperature+Isolate*DeathAge | 97.7 |
| Mature spores load~Temperature*Isolate+DeathAge | 104.8 |
| Mature spores load~Isolate+DeathAge*Temperature | 99.9 |
| Mature spores load~Isolate*DeathAge | 93.2 |
| Mature spores load~Temperature*Isolate | 149.6 |
| Mature spores load~DeathAge*Temperature | 107.8 |
| Mature spores load~Temperature+Isolate+DeathAge | 97.1 |
| **Mature spores load~Isolate+DeathAge** | **92.7** |
| Mature spores load~Temperature+Isolate | 139.0 |
| Mature spores load~DeathAge+Temperature | 104.6 |
| Mature spores load~DeathAge | 100.5 |
| Mature spores load~Isolate | 136.4 |
| Mature spores load~Temperature | 154.8 |
| Mature spores load~1 | 152.4 |

Table S12 : Second cycle cauliflowers

| Step 1 : Random effect selection | |
|---|---|
| Model | AICc |
| ***Second cauliflowers presence~Temperature*Isolate*DeathAge*** | ***205.7*** |
| Second cauliflowers presence~Temperature*Isolate*DeathAge + (1\|Batch) | 206.2 |
| Step 2 : Fixed effect selection | |
| Second cauliflowers presence~Temperature*Isolate*DeathAge | 220.5 |
| Second cauliflowers presence~Temperature+Isolate*DeathAge | 203.8 |
| Second cauliflowers presence~Temperature*Isolate+DeathAge | 205.7 |
| Second cauliflowers presence~Isolate+DeathAge*Temperature | 203.7 |
| Second cauliflowers presence~Isolate*DeathAge | 203.0 |
| Second cauliflowers presence~Temperature*Isolate | 203.6 |
| Second cauliflowers presence~DeathAge*Temperature | 202.7 |
| Second cauliflowers presence~Temperature+Isolate+DeathAge | 199.7 |
| Second cauliflowers presence~Isolate+DeathAge | 198.9 |
| Second cauliflowers presence~Temperature+Isolate | 198.1 |
| Second cauliflowers presence~DeathAge+Temperature | 199.3 |
| Second cauliflowers presence~DeathAge | 198.6 |
| **Second cauliflowers presence~Isolate** | **197.0** |
| Second cauliflowers presence~Temperature | 197.2 |
| Second cauliflowers presence~1 | 196.6 |

Table S13 : Maximal parasitic stage reached

| Step 1 : Random effect selection | |
|---|---|
| Model | AICc |
| **Maximal stage~Temperature+Isolate+DeathAge** | **291.6** |
| Maximal stage~Temperature+Isolate+DeathAge + (1\|Batch) | 293.1 |
| Step 2 : Fixed effect selection | |
| Maximal stage~Temperature*Isolate*DeathAge | NA |
| Maximal stage~Temperature*Isolate+DeathAge | 300.6 |
| Maximal stage~Temperature*DeathAge+Isolate | 293.6 |
| Maximal stage~Temperature+DeathAge*Isolate | NA |
| **Maximal stage~Temperature+Isolate+DeathAge** | **291.6** |
| Maximal stage~Isolate*DeathAge | NA |
| Maximal stage~Temperature*Isolate | 345.7 |
| Maximal stage~DeathAge*Temperature | 296.4 |
| Maximal stage~Temperature+Isolate | 339.3 |
| Maximal stage~Isolate+DeathAge | 299.2 |
| Maximal stage~DeathAge+Temperature | 294.0 |
| Maximal stage~Temperature | 333.6 |
| Maximal stage~Isolate | 341.1 |
| Maximal stage~DeathAge | 300.5 |
| Maximal stage~1 | 335.7 |
